# Substructure of the brain’s Cingulo-Opercular network

**DOI:** 10.1101/2023.10.10.561772

**Authors:** Carolina Badke D’Andrea, Timothy O. Laumann, Dillan J. Newbold, Steven M. Nelson, Ashley N. Nielsen, Roselyne Chauvin, Scott Marek, Deanna J. Greene, Nico U.F. Dosenbach, Evan M. Gordon

## Abstract

The Cingulo-Opercular network (CON) is an executive network of the human brain that regulates actions. CON is composed of many widely distributed cortical regions that are involved in top-down control over both lower-level (i.e., motor) and higher-level (i.e., cognitive) functions, as well as in processing of painful stimuli. Given the topographical and functional heterogeneity of the CON, we investigated whether subnetworks within the CON support separable aspects of action control. Using precision functional mapping (PFM) in 15 participants with > 5 hours of resting state functional connectivity (RSFC) and task data, we identified three anatomically and functionally distinct CON subnetworks within each individual. These three distinct subnetworks were linked to Decisions, Actions, and Feedback (including pain processing), respectively, in convergence with a meta-analytic task database. These Decision, Action and Feedback subnetworks represent pathways by which the brain establishes top-down goals, transforms those goals into actions, implemented as movements, and processes critical action feedback such as pain.

## INTRODUCTION

The human brain is organized into a set of large-scale functional networks that are reproducibly identifiable across populations, datasets, and analysis techniques ^1–7^. One of the most functionally complex large-scale networks is the Cingulo-Opercular network (CON). The CON is composed of a set of functionally coupled regions in dorsal anterior cingulate cortex (dACC), dorsomedial prefrontal cortex (dmPFC), anterior insula (aI), supramarginal gyrus (SMG), pars marginalis of the cingulate gyrus, and the anterior prefrontal cortex (aPFC) ^5,6,8–10^. The CON exhibits strong functional connectivity to a highly diverse set of brain networks, including cognitive, sensory, and somatomotor networks ^2,5,6^. This intrinsic connectivity structure suggests that CON may exert influence over a wide variety of brain processes, from high-level cognition to basic motor outputs.

The CON has been primarily characterized as a network critical for exerting top-down cognitive control over other purely cognitive functions. Large signals are observed in the CON when complex cognitive tasks are initiated, and it also exhibits sustained task signals to maintain goals and prevent distraction ^8–10^. CON regions also respond strongly to feedback in order to provide more effective top-down control in the future. The CON exhibits large responses when errors are made or when reaction times are slow, and CON signals are elevated when stimuli are ambiguous or multiple possible responses are in conflict ^8–19^.

Other lines of work suggest that some CON regions enable top-down motor control ^20^. Recent work has demonstrated direct connections between the CON and Somato-cognitive action network (SCAN) regions that alternate with effector-specific regions in primary motor cortex (M1); these connections are thought to represent a mechanism by which the CON implements whole-body action plans within the motor system ^21^. Further, prolonged dominant arm immobilization results in both substantial changes in motor behavior and strengthened functional connectivity between disused M1 and the CON, suggesting an important role in motor control ^22,23^. In non-human primates, three cingulate areas have been identified as critical for motor planning (termed the rostral, ventral, and dorsal cingulate motor areas) ^24,25^; their human homologues (anterior/posterior rostral cingulate zone, and caudal cingulate zone) are likely located within the anterior cingulate portion of the CON.

The CON also plays a role in processing painful stimuli. The dACC and the aI are commonly reported as brain regions most active during application of painful stimuli ^26–28^. This pattern is generally consistent across both somatic and visceral pain ^29^, is spatially distinct from representations of negative affect or social pain ^28^, and partially overlaps with CON regions involved in cognitive control ^30^.

The substantial functional heterogeneity exhibited by the CON, including control over cognitive processes, control over action plans, and processing of painful stimuli, suggests that the CON is not well represented as a single brain system with a unitary label. Instead, it is likely that the apparently unitary CON is a representation of network structure at just one level of a functional hierarchy ^31–35^ that could be further divided into functionally divergent substructures. Indeed, recent work has suggested at least a bipartite division of CON, with one CON subdivision linked to top-down cognitive control and another linked to motor and interoceptive functions ^36^, though these divisions were identified using a priori ROIs in group-averaged data, and so are necessarily imprecise.

To improve precision of network delineation, our group has developed a technique called precision functional mapping (PFM) that uses repeated fMRI scanning to reliably characterize brain networks within individual highly sampled participants ^2,36–47^. By characterizing networks at the individual level, detailed subdivisions within large-scale networks have been identified, including within the DMN^35,48^, striatum^49^, and primary motor cortex ^21^.

Here, we implemented a similar approach in 15 participants with > 5 hours of RSFC data and additional task data, in order to identify and characterize discrete substructures within the CON that could explain the organization and integration of various functional domains. We hypothesized that multiple distinct subnetworks would be identified in each participant that would be specialized for controlling complex cognitive functions and adjudicating among multiple options, developing and controlling motor actions, and processing pain-related and other feedback.

## RESULTS

### Three Distinct CON Subnetworks in Each Individual

The subnetwork structure of each individual’s CON was identified following previously described methods using the data-driven Infomap community detection algorithm ^21,35,49^. Three spatially distinct subnetworks were identified within the large-scale CON. An example participant is shown in Figure 1. These three subnetworks were topologically consistent across participants (Figure 2). All three subnetworks were identified in each of the 15 PFM data sets (Figure S1). Subnetworks were initially labeled according to their locations on the cortex as the Anterior, Central, and Lateral subnetworks of CON. Together, these subnetworks included representations in each of the major areas of the CON, including: 1) the dorsal anterior cingulate, dorsal anterior insula, and anterior PFC (Anterior); 2) the posterior dorsomedial prefrontal cortex, inferior supramarginal gyrus, posterior middle insula, and dorsal postcentral gyrus (Central); 3) the anterior middle insula, anterior lateral frontal cortex, inferior frontal gyrus, dorsal supramarginal gyrus, and pars marginalis of the cingulate (Lateral).

**Figure 1:**
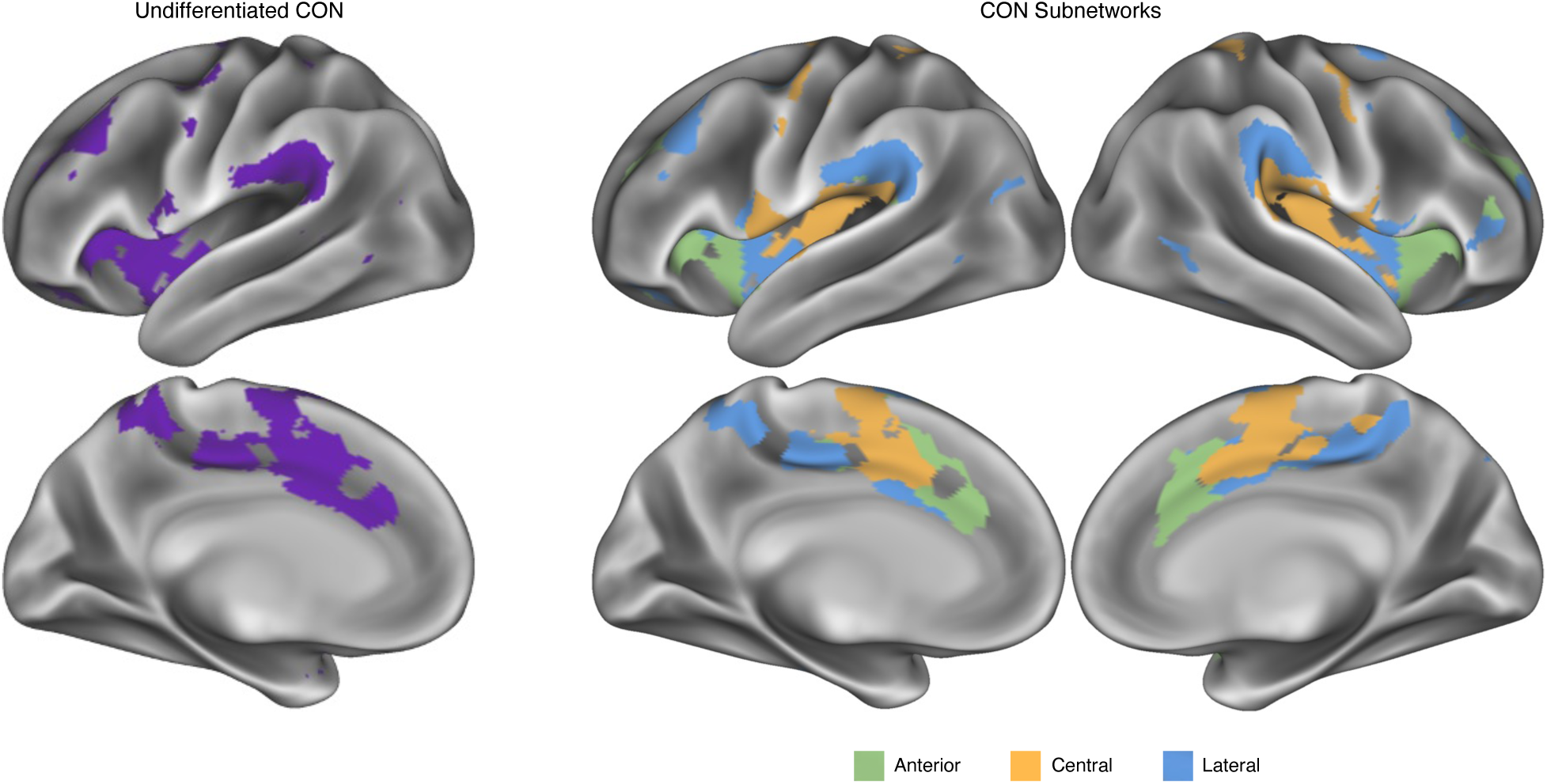
Three Cingulo-Opercular Subnetworks. The undifferentiated CON (left) and CON subnetworks (right) in an exemplar participant (P01) with 356 minutes of resting-state fMRI data. Three distinct subnetworks were consistently identified in every individual: an Anterior subnetwork in dorsal anterior cingulate, dorsal anterior insula, and anterior PFC (green); a Central subnetwork in more posterior dorsal anterior cingulate extending up to dmPFC, middle/posterior insula, and regions just anterior and posterior of the central sulcus (yellow); and a Lateral subnetwork in middle insula, supramarginal gyrus, and the pars marginalis of the cingulate (blue). See Figure S1 for all individual participants.

**Figure 2:**
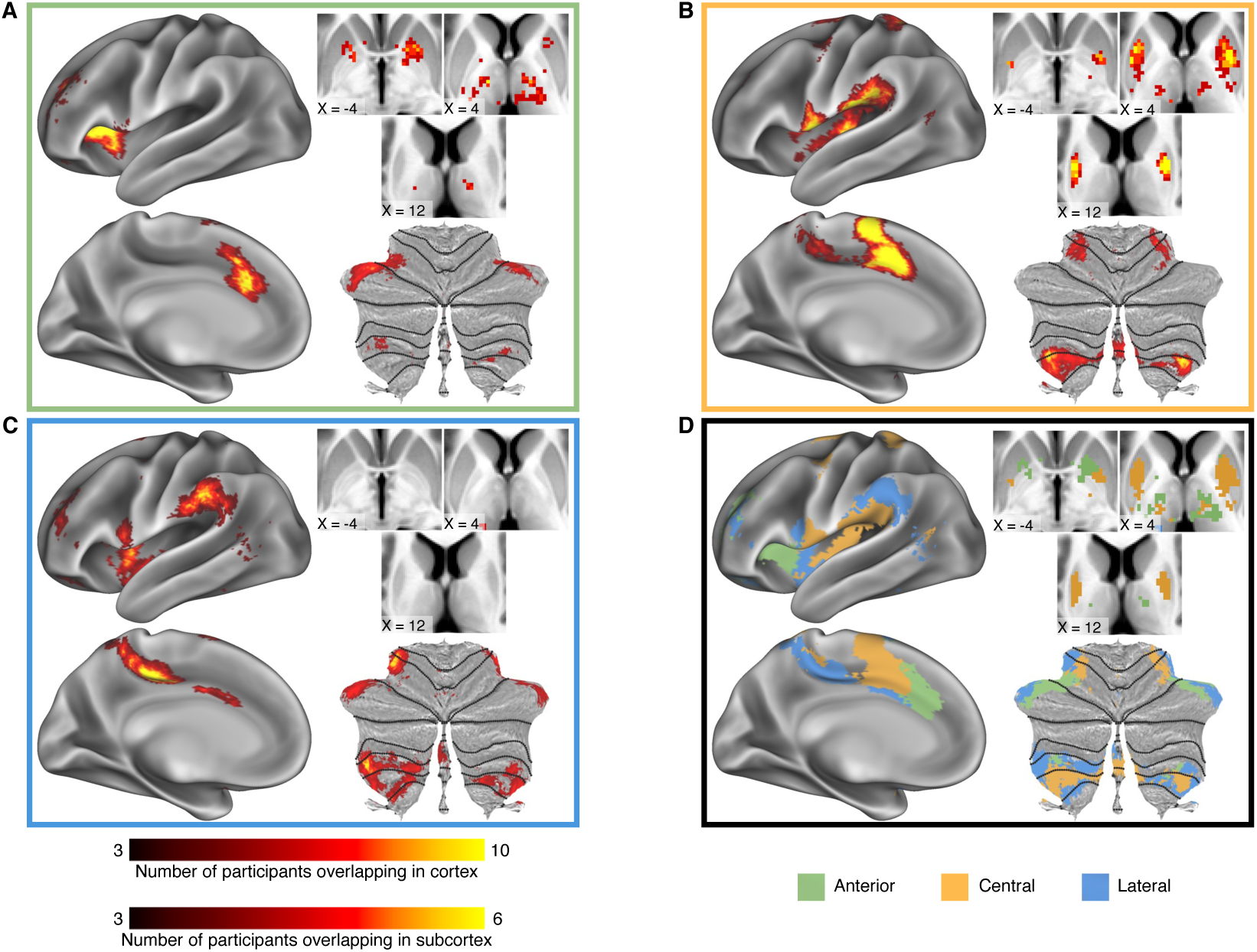
Cortical and subcortical distribution of CON subnetworks across participants. A-C) Density maps illustrate the number of participants with overlapping subnetwork representations at each point in cortex (left), thalamus and striatum (top right), and cerebellum (bottom right), for the A) Anterior, B) Central, and C) Lateral subnetworks. Maps were thresholded to retain points at which at least three participants exhibited overlap. Differential scaling of density maps in cortex and subcortex was employed because of increased cross-participant variability (due to lower SNR) in subcortex. D) A winner-take-all map illustrates the relative topographies of all three subnetworks.

The Anterior and Central subnetworks had representation in ventral and dorsal anterior putamen, respectively, though the Central subnetwork had more extensive putamen representation, and in central thalamus (Fig 2A-B). By contrast, almost no voxels were consistently identified as part of the Lateral subnetwork within striatum or thalamus. All three subnetworks were represented in the cerebellum, though overlap across participants was lower in cerebellum than in cortex or striatum, suggesting greater anatomical heterogeneity across individuals.

### CON subnetworks are functionally dissociated by Meta-Analytic Network Annotation

The functional role of each CON subnetwork was determined by matching the subnetwork to distributed activation patterns within the Neurosynth meta-analysis database ^50^. We term this novel connectome annotation method Meta-Analytic Network Annotation.

For each CON subnetwork, we first identified all articles in the database that reported multiple activation peaks that were at least partially congruent with the spatial distribution of that subnetwork (see Methods). Then, for each of the 742 cognition-related terms in the database, a one-way ANOVA tested whether the weighting of that term within matched articles was significantly different across the three subnetworks. Each term with significant differences across the subnetworks was associated with the subnetwork for which it had the highest weighting. Word clouds illustrate the terms associated with each subnetwork, with word size scaled to the magnitude of differences among subnetworks (Figure 3). Terms shown in black exhibited differences between subnetworks that were significant at p < 0.05 (unc.); colored terms were significant at p < 0.05, FDR corrected for the total number of terms tested. It should be noted that while this analysis identifies which subnetwork is most strongly associated with each term, such results should not be misinterpreted as an absence of association for other subnetworks. For example, while the Lateral subnetwork is most strongly associated with “pain”, the Anterior and Central subnetworks were also pain associated, though less strongly.

**Figure 3:**
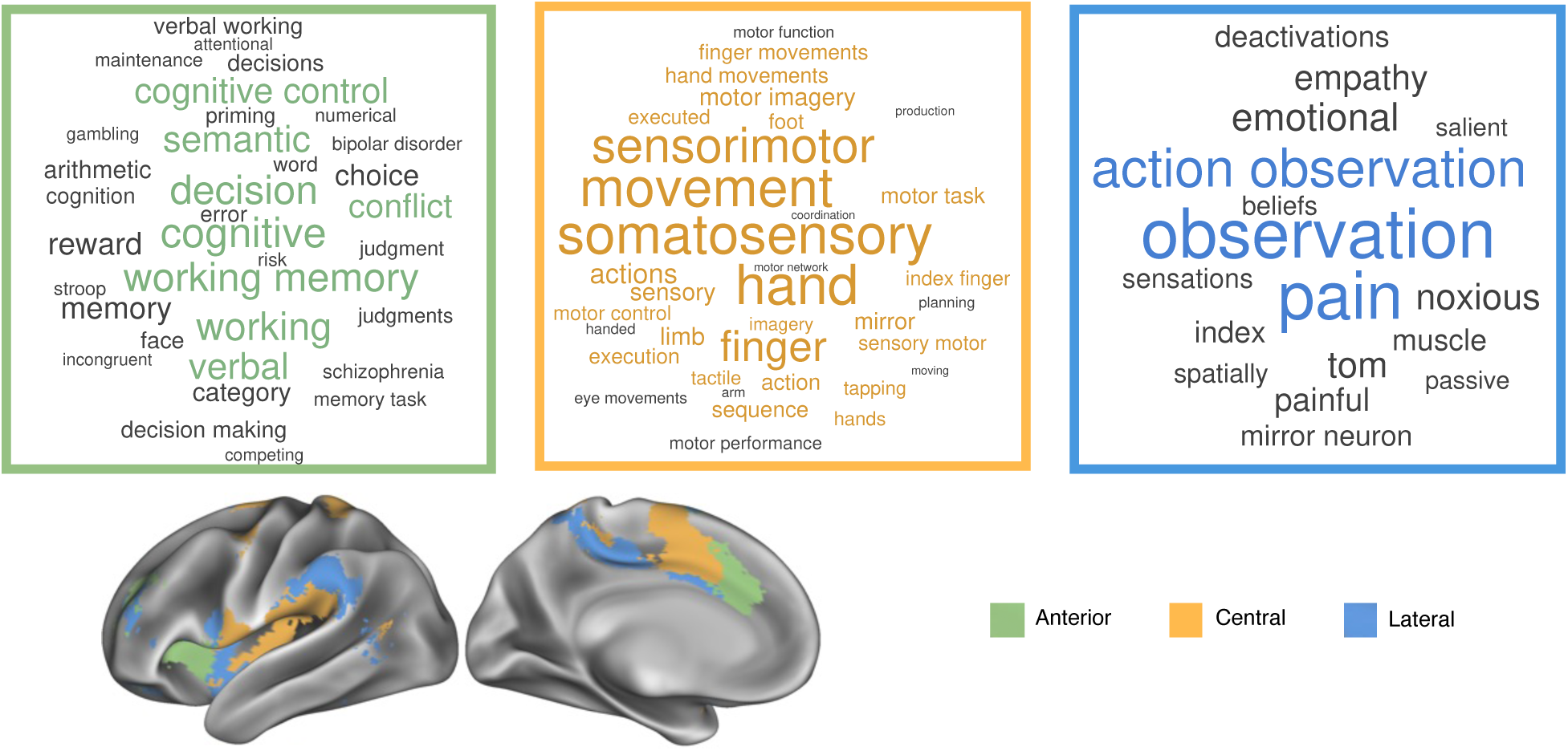
Meta-Analytic Network Annotation of Cingulo-Opercular subnetworks. Subnetworks from the cross-participants winner-take-all analysis (Fig 2D) were matched to spatial activation distributions that had task descriptor terms in the Neurosynth database ^50^. Word clouds illustrate terms more associated with activation patterns of each Cingulo-Opercular subnetwork compared to the other subnetworks (tested for each term via one-way ANOVA). Larger font size indicates higher frequency of the term. Terms shown in black are significant at p < 0.05 (unc.); colored terms are significant at p < 0.05, FDR corrected for the 742 terms tested.

The Anterior subnetwork was significantly more associated with terms such as ‘decision’, ‘cognitive control’, ‘conflict’, ‘semantic’, and ‘working memory’ than the other subnetworks. It was also most associated with terms such as ‘reward’, ‘risk’, and ‘error’, though these tests did not survive correction for multiple comparisons. Together, these terms suggest a circuit that integrates information from multiple sources to decide on a course of action. Thus, we functionally annotate the Anterior CON as the Decision subnetwork.

The Central subnetwork was associated with motor terms; it included pure motor terms such as ‘movement’, ‘hand’, and ‘finger’, as well as motor control terms such as ‘actions’, ‘motor control’, ‘execution’, and ‘motor imagery’. It also included sensory feedback terms such as ‘somatosensory’. These terms suggest a circuit that generates an action plan and receives sensory feedback about that action. We annotate this as the Action subnetwork.

The Lateral subnetwork was associated with terms such as ‘pain’ and ‘noxious’, but also with terms such as ‘action observation’. These terms suggest a circuit that processes multiple somatic, non-cognitive feedback modalities resulting from actions, such as visual feedback and pain. We annotate this as the Feedback subnetwork.

### CON subnetworks exhibit differential connectivity with other large-scale networks

Spring-embedded graph visualizations (following methods from ^5,21,53^; example participant shown in Figure 4A; all participants in Figure S2) suggest that the three CON subnetworks exhibit preferential connectivity to distinct large-scale networks. Quantification of average connectivity to each network showed that the Action CON subnetwork was most closely linked to motor networks, the Decision CON subnetwork was most closely linked to the Salience network, and the Feedback CON subnetwork was most closely linked to the Dorsal Attention Networks (Figure 4B).

**Figure 4:**
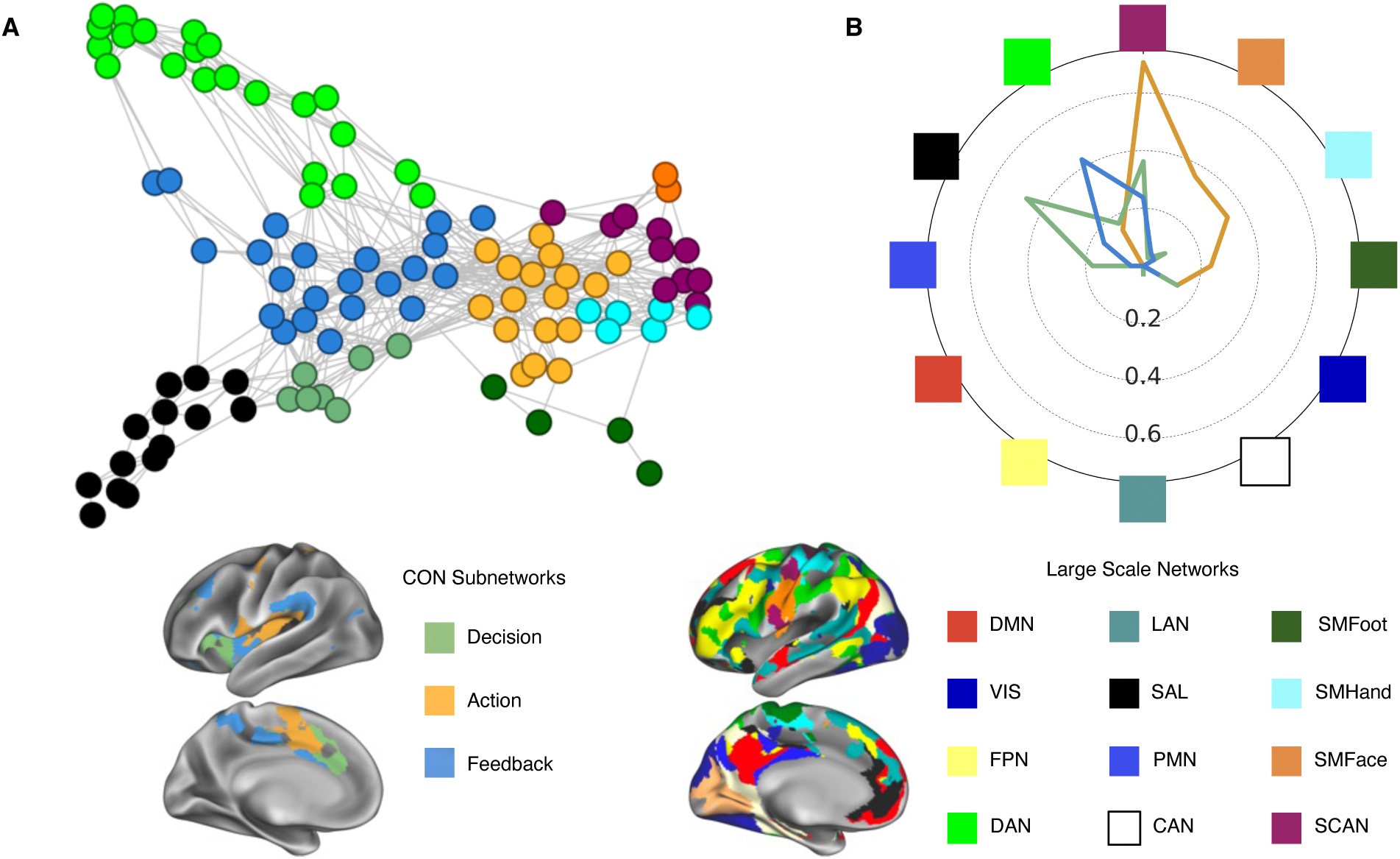
Functional connectivity of CON subnetworks. A) A spring embedded plot in an exemplar participant (P01) illustrates the preferential connectivity of Cingulo-Opercular subnetworks to large-scale functional networks. For clarity of visualization, only networks most closely associated with the subnetworks are shown. See Figure S2 for all individual participants. B) Across participants, individual-specific Cingulo-Opercular subnetworks demonstrate preferential connectivity to other individual-specific large-scale networks. The radial axis indicates the strength of functional connectivity Z(r) between each CON subnetwork and each large-scale functional network. Negative connectivity values are not represented. Colors and spatial locations of CON subnetworks (left) and other large-scale networks (right) are shown in the exemplar participant at the bottom.

Paired t-tests were used to statistically compare the strength of functional connectivity between pains of CON subnetworks to a given large-scale network. Reported findings all survived FDR correction for the number of comparisons run at q < 0.05. Across participants, the Decision subnetwork exhibited stronger connectivity with the Salience Network than did any other subnetwork (ts(14) > 2.9, ps(unc) < 0.012). The Feedback subnetwork exhibited stronger connectivity with the Dorsal Attention Network (ts(14) > 3.2, ps(unc) < 0.007) than any other network. The Action subnetwork exhibited stronger connectivity with SCAN (ts(14) > 5.8, ps(unc) < 0.001) as well as Somatomotor Foot, Hand, and Face networks (ts(14) > 5.4, ps(unc) < 0.001). The Action subnetwork also exhibited significantly stronger negative connectivity with the Default network compared to the other two subnetworks (ts(14) > 3.1, ps(unc) < 0.008). All subnetwork-network t-tests can be found in Supplementary Table 1.

### Rs-fMRI signals from CON subnetworks are temporally ordered

Previously, we have shown that low-frequency rs-fMRI signals exhibit different timings within different subnetworks of the Default Mode and action and motor networks ^21,35^. Here, lag analyses demonstrate the temporal order of low-frequency BOLD signals within networks of the putative action output processing stream, including CON subnetworks, SCAN, and effector-specific hand motor networks (Figure 5). A one-way ANOVA found a significant main effect of subnetwork/network identity (F(4,74) = 5.49, p = 0.0007). Post-hoc t-tests indicated that signals in the Feedback subnetwork occurred later than those in the Action, SCAN, and Somatomotor Hand networks (ts(14) > 2.62, ps < 0.02). Signals in the Decision subnetwork occurred later than those in the Somatomotor Hand (ts(14) > 2.44, p = 0.02), as did signals in the Action subnetwork, though only at trend level (t(14) = 1.76, p = 0.09). All temporal lag t-tests can be found in Supplementary Table 2. Previous reports have associated inter-regional lags in infra-slow (<0.1 Hz) signals with propagation of higher-frequency delta activity (0.5-4 Hz) in the opposite direction ^52^. Thus, these results suggest that high-frequency activity propagates from Feedback and Decision subnetworks to the Action subnetwork, and then to SCAN and effector-specific Somatomotor networks.

**Figure 5:**
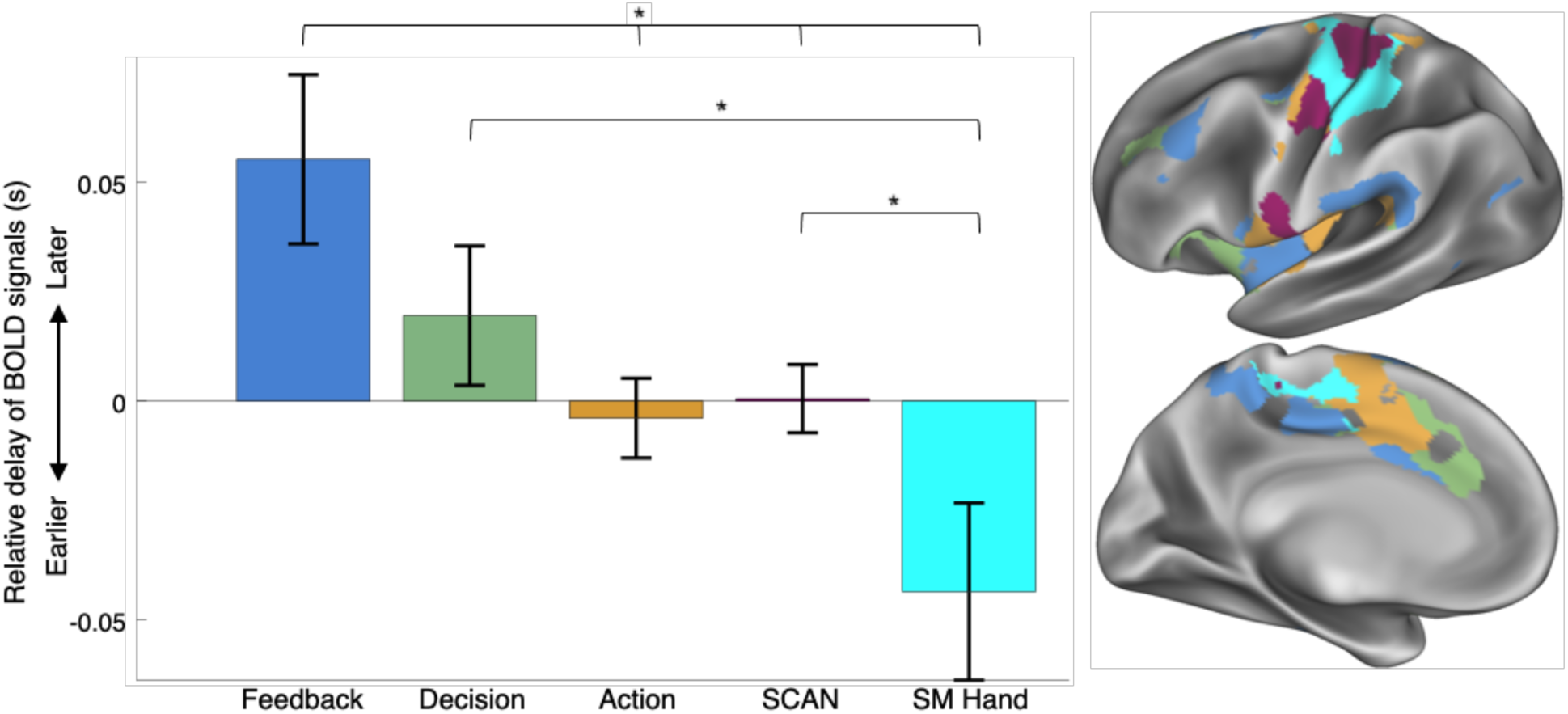
Temporal ordering of CON signals in the action output hierarchy. Temporal ordering of subnetwork fMRI signals for each individual-specific CON subnetwork, as well as for the SCAN and the Somatomotor-hand networks. Values are averaged across vertices within each subnetwork and across participants. Standard error across participants is indicated by error bars. A one-way ANOVA indicated a significant main effect of subnetwork/network identity (p = 0.0007). * indicates p < 0.05 for post-hoc paired t-tests. Inset shows subnetwork and network topography for an example participant (P01). Prior electrophysiology work suggests that later infra-slow activity (here, the Feedback subnetwork) corresponds to earlier delta-band (0.5-4Hz) activity ^52^.

### CON subnetworks exhibit differential evoked responses during cognitive and motor tasks

Task data collected in 10 of the 15 participants across five motor task conditions (HCP motor task ^53^: Tongue, Left Hand, Right Hand, Left Leg, Right Leg) and two cognitive conditions (spatial discrimination and verbal discrimination ^2^) confirmed differences in evoked activity between CON subnetworks.

Paired t-tests (FDR-corrected for all comparisons run) revealed that CON subnetworks were differentially activated across the seven task conditions (Figure 6). In the Tongue movement condition, the Decision and Action subnetworks were more activated than the Feedback subnetwork (ts(9) > 3.56, ps(unc) < 0.007). For all Hand and Leg movement conditions, the Action subnetwork was more activated than either of the Decision or Feedback subnetworks (ts(9) > 3.80, ps(unc) < 0.005). For each of the two cognitive conditions (spatial and verbal discrimination), the Decision subnetwork was more active than either of the other two subnetworks (ts(9) > 4.42, ps(unc) < 0.002). These results demonstrate that CON subnetworks are distinguished by within-participant differences in task activation that are consistent with the Neurosynth-based Meta-Analytic Network Annotation. Specifically, the Action CON subnetwork is more active during motor conditions, and the Decision CON subnetwork is more active during this particular set of cognitive conditions. The Feedback subnetwork was not strongly activated during any condition, suggesting that this specific set of task conditions was not well-suited to highlight its function. T-tests for all subnetworks, for all tasks, can be found in Supplementary Table 3.

**Figure 6:**
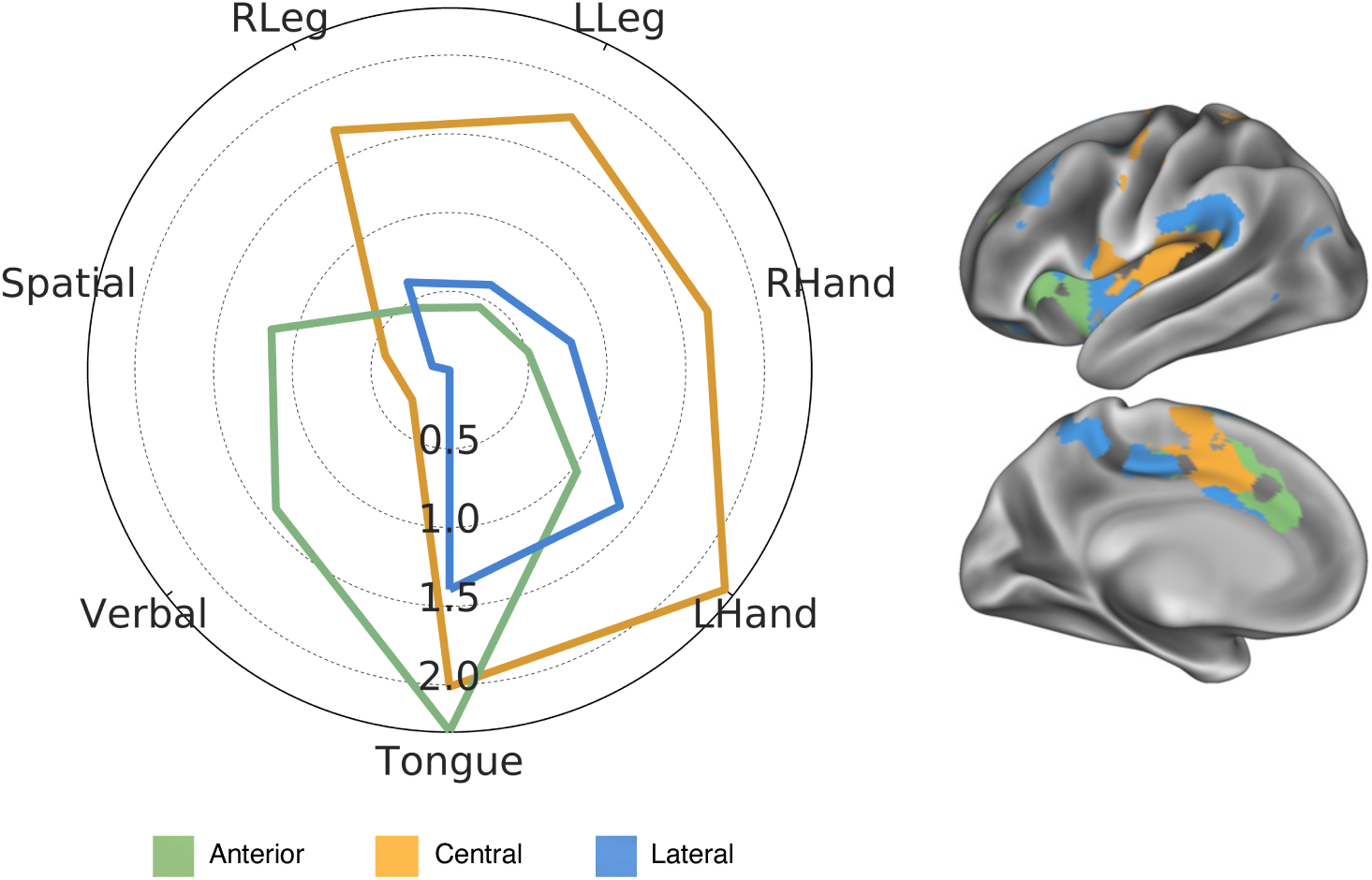
Differentiation of Cingulo-Opercular subnetworks based on task activations. Ten participants performed motor tasks^53^ (including flexure of right and left toes, open/closing of right and left hand, and left-right movement of the tongue), as well as a Spatial and a Verbal Discrimination task^54^. The radial axis indicates the z-score of each condition relative to baseline fixation, averaged across all participants and across all vertices in each individual-specific CON subnetwork. Inset shows subnetwork topography for an example participant (P01).

## DISCUSSION

We identified three distinct subnetworks within the large-scale Cingulo-Opercular network. Broadly, these subnetworks appear to support the various known functions of the CON. Based on the connections, meta-analytic network annotation, and prior descriptions of each subnetwork’s constituent regions in the literature, we argue that these three subnetworks can be conceptualized as a Decision subnetwork, an Action subnetwork, and a Feedback subnetwork. Together, identification of these network substructures helps inform how the human brain develops goals and implements actions to serve those goals.

### A Decision subnetwork for action selection, maintenance, and control

The Decision subnetwork described here converges closely with the most well-described functions of the classic CON. The functional core of the classic description of CON ^8–10^ is centered in bilateral dACC, aI, and aPFC, which exactly converges with the Decision subnetwork described here. The preferentially cognitive nature of functions associated with the Decision subnetwork here also converges with previous reports of CON functionality, including involvement in task maintenance, error processing, conflict monitoring, and ambiguity processing to decide among competing options ^9,14,17–19,55^. The Decision subnetwork had representation in the anterior ventral putamen, convergent with dACC-ventral anterior putamen projections in non-human primates argued to be related to selecting among different actions^56^.

Notably, the Decision subnetwork was most strongly connected to the Salience network ^57^. Salience network regions have been associated with reward valuation and motivation in meta-analyses ^58^, and dorsal anterior cingulate regions corresponding with the Decision subnetwork have been argued to evaluate reward information in order to make decisions ^59^, guide behavior ^60,61^, and produce actions ^62^. This suggests that the Salience network inputs to the Decision subnetwork enable it to select decisions, exert control, and produce actions in ways that are guided by the reinforcement value of that decision. The outputs of the Decision subnetwork may be abstract—exerting control over working memory or attention functions—or they may be physical actions. In the latter case, the temporal ordering of signals (Fig 5) suggests that Decision network outputs are projected to the Action subnetwork.

### An Action subnetwork for initiation and top-down control over physical actions

Action control can be understood as a cascade of executive functions that proceed from abstract to concrete action plans, and then to execution of movement ^63^. The CON in particular has been shown to exhibit strong connectivity to SCAN in the precentral gyrus ^21^ and to exert influence over motor functions ^23,64^. Further, non-human primate research has demonstrated that the regions at the top of this motor control hierarchy are the rostral and caudal cingulate motor zones in the dorsal medial prefrontal cortex ^24,25^, which initiate a chain of neuronal projections backwards to supplementary motor area, premotor area, and finally to primary motor cortex. These cingulate motor zones appear homologous with the dorsomedial prefrontal portion of the CON, and they are most likely identified here as the Action subnetwork, which exhibited strong connectivity with motor networks, and strong activation—in both participant-specific tasks and in Meta Analytic Network Annotion—during motor function.

The Action subnetwork was particularly strongly connected to the recently identified SCAN ^21^, which has been argued to implement action control on a whole-body level, including regulation of posture and internal physiology (e.g., blood pressure, adrenaline release), to maintain body allostasis. The action subnetwork had substantial representation in dorsal anterior putamen, which is the potion of striatum associated with motor planning and learning, and receiving projections from premotor areas in non-human primates^56^. Further, several Action subnetwork regions were directly proximal to either primary somatosensory cortex (S1) on the postcentral gyrus, or secondary somatosensory cortex (S2) in the dorsal posterior insula. Further, meta-analytic network annotation indicated somatosensory functions in the Action subnetwork. Thus, the Action subnetwork may also receive fast sensory feedback from somatosensory functions about the immediate results of executed actions. Given these postulated functions, it is possible that the Action subnetwork also represents visceral sensation, which has localization in middle insula convergent with this subnetwork^65^.

### A Feedback subnetwork for evaluating action outcomes

Unlike the Decision and Action subnetworks, the Feedback subnetwork is notable in exhibiting a paucity of striatal or thalamic representation. Cortico-striato-thalamo-cortical loops are argued to be critical for the motivation, development, planning, and execution of goal-directed actions ^56^. The absence of such loops in the Feedback subnetwork suggests that this circuit is not directly involved in the feedforward hierarchy of action production and control.

Instead, we hypothesize that the Feedback subnetwork provides more temporally extended feedback to the Decision subnetwork to evaluate the efficacy of generated plans. This subnetwork did not strongly activate during performance of either motor or cognitive tasks, but was strongly associated with meta-analytic terms reflecting processing of painful and noxious stimuli. Further, the subnetwork’s representation in the middle section of the anterior insula is consistent with known distributions of pain-related brain activity ^27,29^. Nociceptive somatosensory stimulation has been shown to increase functional connectivity between CON, sensorimotor, and emotional networks. This suggests a role for the CON, and specifically the Feedback subnetwork, in attending to, integrating, and transmitting sensory information related to pain ^66^. Notably, cortical processing of pain information may operate on a different timescale than the immediate spinal-mediated reflexive responses to pain, reflecting a slower, more cognitive interpretation of pain.

However, the Feedback subnetwork cannot be conceptualized as purely pain-related. In addition to pain, the Feedback subnetwork was also associated with meta-analytic terms reflecting observation of actions; and further, it exhibited uniquely strong connectivity to the Dorsal Attention Network, which helps direct eye movements and processes attention-directed visuospatial information ^67,68^. Together, we hypothesize that the Feedback subnetwork processes multiple types of post-action feedback, including both pain and visual information about the results of an executed action. Most interestingly, Feedback subnetwork-associated terms included “empathy”, “tom [theory of mind]”, and “mirror neuron,” suggesting this system may also process painful and visual feedback information about the results of actions taken by others.

The Feedback subnetwork may not be as primary in processing another type of feedback, task errors (i.e. incorrect choices). Here, the “Error” term was associated most with the Decision subnetwork in meta-analytic network annotation, though this difference only passed an uncorrected threshold. Prior work has shown that the strongest error-related signals are in Decision subnetwork regions such as dACC and aI, but weaker error signals can also observed in some Feedback regions such as inferior parietal and inferior frontal gyri ^8,13,14^. We hypothesize that task errors, being more complex and context-dependent than pain or visual feedback, may require more integrated cross-subnetwork processing, and ultimately adjudication by the Decision subnetwork.

### Information flow through the action hierarchy

Differences in rs-fMRI relative signal timings among the subnetworks suggest that information primarily flows from Feedback to Decision to Action subnetworks, and then to the SCAN and the motor system and likely back to Feedback completing the cycle. Infra-slow rs-fMRI signals were detected relatively later in Feedback and Decision CON subnetworks than in motor systems (SCAN and effector-specific). However, the “sender-receiver” model argues that infra-slow activity (<0.1 Hz) propagates in the opposite direction from the higher frequency (i.e. delta-band, 0.5-4 Hz) cortical activity reflective of neural processing, in order to coordinate the timing of high-frequency information exchange via phase-amplitude coupling ^52,69^. This result expands findings from our prior work, which described a CON->SCAN->effector-specific M1 ordering of signals ^21^, but assumed homogeneity of CON itself. Here, the distinct ordering of signals we observed propagating within and beyond CON converge with prior work conducting invasive recordings in macaques, which found that signals in more rostral cingulate regions occur much earlier before a movement than those in caudal cingulate ^70^.

### A hierarchy of goal establishment and action implementation

Together, these findings suggest a hierarchical model of goal and action control in the brain (Figure 7). In this model, the Decision subnetwork receives connections reflecting valuations from regions in the Salience network that process reward and incorporates this information to perform judgments, adjudicate between alternatives, and initiate and maintain a goal. If the goal can be expressed as a physical action, it is projected backwards to the Action subnetwork where a series of operations transform the abstract goal into a concrete action plan. This action plan is projected to the Somato-Cognitive Action Network to prepare the whole body to implement the plan ^21^, and finally to effector-specific M1 where it is transformed into movement. Feedback about the action’s outcomes (including salient visual stimuli and pain) is provided through the Feedback subnetwork to the Decision subnetwork, which processes these signals, as well as goal-incompatible errors, to modify maintained goals and future planned actions. Delineation of this functionally heterogeneous network substructure has critical implications for the characterization of neurological and psychiatric patients with disruptions along the hierarchy of action control, including ADHD, abulia, OCD, and Parkinson’s disease. Further, the ability to specifically identify this individually variable subnetwork structure in every single individual could set the stage for important advances in personalized, precision-targeted neuromodulatory treatments for these disorders.

**Figure 7:**
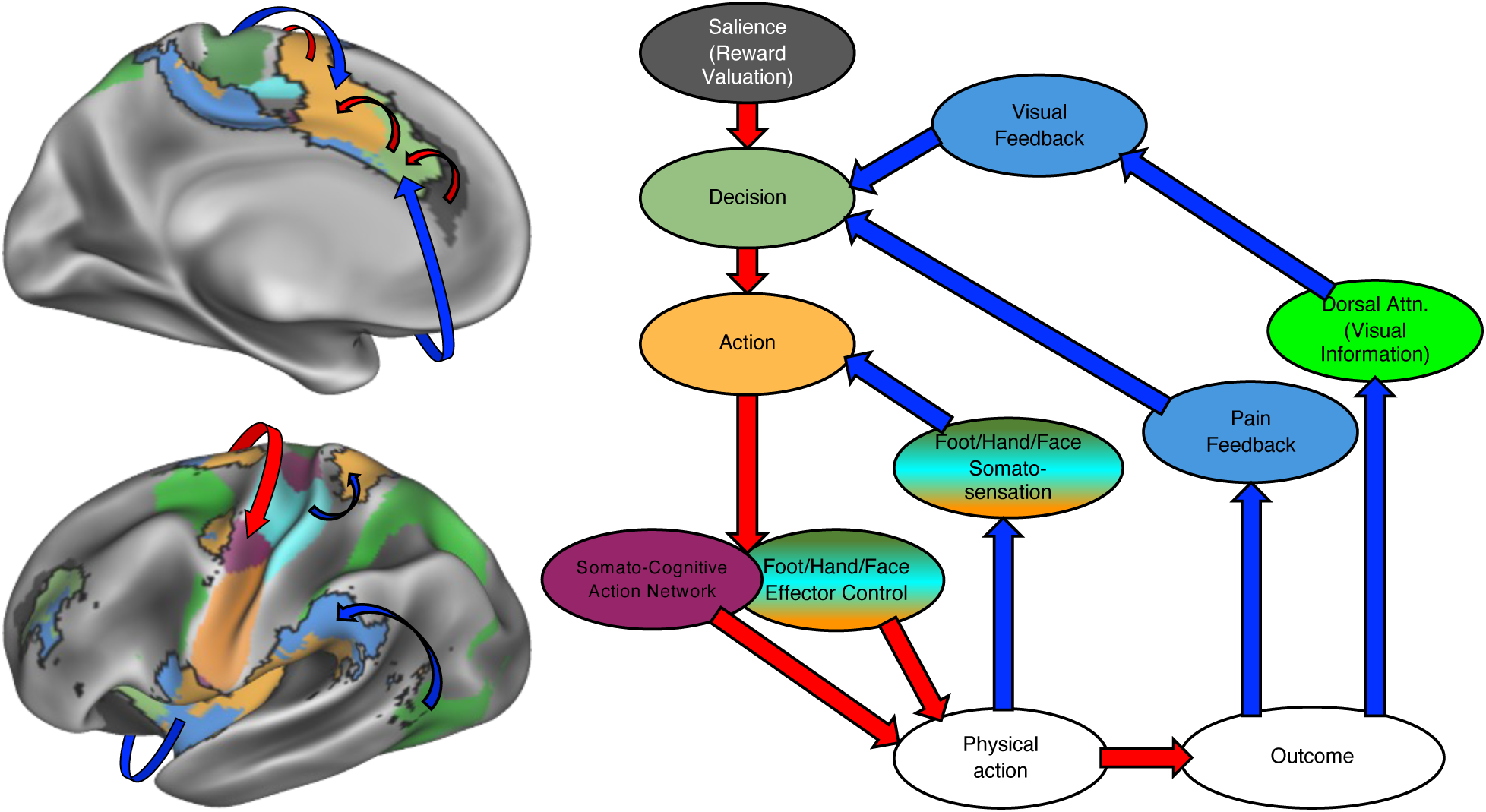
Model of CON subnetworks within a hierarchical organization of goal and action control. Our proposed model represented on the cortex (left) and in schematic form (right). Reward-driven valuations from the Salience network feed into CON subnetworks, which establish goals (Decision subnetwork) and goal-related action plans (Action subnetwork), and project those action plans to the SCAN and to effector-specific M1 to execute an action. Fast somatosensory information resulting from the action is processed by the Action subnetwork, which modifies action plans. Slower goal-related visual and painful outcomes of the action are processed by the Feedback subnetwork, which modifies goals represented by the Decision subnetwork. Red arrows indicate feedforward projections; blue arrows indicate feedback.

## ACKNOWLEDGEMENTS

This work was supported by NIH grants NS110332 (DJN), MH121518 (SM), MH129616 (TOL), MH096773 (NUFD), MH122066 (EMG, NUFD), MH121276 (EMG, NUFD), MH124567 (EMG, NUFD), NS129521 (EMG, NUFD), and NS088590 (NUFD); by the National Spasmodic Dysphonia Association (EMG); by the Taylor Family Foundation (TOL); by the Intellectual and Developmental Disabilities Research Center (DJG, NUFD); by the Kiwanis Foundation (NUFD); by the Washington University Hope Center for Neurological Disorders (EMG, NUFD); and by Mallinckrodt Institute of Radiology pilot funding (DJG, EMG, NUFD).

## CONTRIBUTIONS

Conceptualization: C.B.D., E.M.G.,

Data curation: D.J.N., N.U.F.D., E.M.G.

Formal analysis: C.B.D., E.M.G.

Supervision: E.M.G.

Writing – Original Draft: C.B.D., E.M.G.

Writing – Review & Editing: All authors

## METHODS

Data for this project was compiled from three preexisting datasets.

### Dataset 1: Plasticity dataset

#### Participants

Data were collected from 3 healthy, right-handed adult participants (ages 35, 25, and 27; one female; identified here as P01-P03) as part of a study investigating effects of arm immobilization on brain plasticity (data previously published in ^21–23,71^. Two participants are authors (NUFD and ANN). The remaining participant (male, 27 years old) was recruited from the Washington University community. Informed consent was obtained from all participants. The study was approved by the Washington University School of Medicine Human Studies Committee and Institutional Review Board. All data employed here was collected either prior to the immobilization intervention (Participants P01-03) or two years afterwards (Participant P02), and so we do not report details of that intervention.

#### MRI image acquisition

Participants were scanned every day for twelve consecutive days using a Siemens Prisma 3T scanner on the Washington University Medical Campus. Every session included a 30-minute resting-state fMRI scan collected as a blood oxygen level-dependent (BOLD) contrast sensitive gradient echo-planar sequence (TE=33 ms, flip angle=84°, resolution=2.6 mm isotropic, TR=1100 ms, multiband 4 acceleration). During this scan, participants were instructed to hold still and look at a white fixation crosshair presented on a black background. A pair of spin echo EPI images with opposite phase encoding directions (AP and PA) but identical geometrical parameters to the BOLD data were acquired to correct spatial distortions.

For all fMRI data, head motion was tracked in real time using Framewise Integrated Real-time MRI Monitoring software (FIRMM ^72^). An eye-tracking camera (EyeLink, Ottawa) was used to monitor participants for drowsiness.

For Participant 01 and 03, every session also included collection of a high-resolution T1-weighted MP-RAGE (TE=2.22ms, TR=2400ms, flip angle=8°, 208 slices with 0.8×0.8×0.8mm voxels) and a T2-weighted spin-echo image (TE=563ms, TR=3200ms, flip angle=120°, 208 slices with 0.8×0.8×0.8mm voxels).

For Participant 02, the structural images used were collected as part of Dataset 2 and consisted of four T1-weighted images (sagittal, 224 slices, 0.8 mm isotropic resolution, TE=3.74 ms, TR=2400 ms, TI=1000 ms, flip angle = 8°) and four T2-weighted images (sagittal, 224 slices, 0.8 mm isotropic resolution, TE=479 ms, TR=3200 ms) collected on a Siemens TRIO 3T scanner.

### Dataset 2: Midnight Scan Club Dataset

#### Participants

Data were collected from 10 healthy, right-handed, young adult participants (5 females; age: 24–34; identified here as P08-P15). The participants were recruited from the Washington University community. Other findings using these participants have been previously reported in ^2,40,45,46,73^. Two participants were excluded because they were also participants in the Plasticity dataset. One participant is an author (SMN). Informed consent was obtained from all participants. The study was approved by the Washington University School of Medicine Human Studies Committee and Institutional Review Board.

#### MRI image acquisition

Imaging for each participant was performed on a Siemens TRIO 3T MRI scanner over the course of 12 sessions conducted on separate days, each beginning at midnight. Structural MRI was conducted across two separate days. In total, four T1-weighted images (sagittal, 224 slices, 0.8 mm isotropic resolution, TE=3.74 ms, TR=2400 ms, TI=1000 ms, flip angle = 8 degrees), four T2-weighted images (sagittal, 224 slices, 0.8 mm isotropic resolution, TE=479 ms, TR=3200 ms), four MRA (transverse, 0.6 x 0.6, x 1.0mm, 44 slices, TR=25ms, TE=3.34ms) and eight MRVs, including four in coronal and four in sagittal orientations (sagittal: 0.8 x 0.8 x 2.0 mm thickness, 120 slices, TR=27 ms, TE=7.05ms; coronal: 0.7 x 0.7 x 2.5 mm thickness, 128 slices, TR=28ms TE=7.18 ms), were obtained for each participant. Analyses of the MRA and MRV scans are not reported here. On ten subsequent days, each participant underwent 1.5 hours of functional MRI scanning beginning at midnight. In each session, we first collected thirty contiguous minutes of resting state fMRI data, in which participants visually fixated on a white crosshair presented against a black background. Each participant was then scanned during performance of three separate tasks: motor (2 runs per session, 7.8 minutes combined); mixed design including both a spatial and a verbal discrimination condition (2 runs per session, 14.2 minutes combined); and incidental memory (3 runs per session, 13.1 minutes combined). Preliminary analysis of the incidental memory task indicated that CON subnetworks were not active during this task, so we do not report results in detail here. Across all sessions, each participant was scanned for 300 total minutes during the resting state and approximately 350 total minutes during task performance. All functional imaging was performed using a gradient-echo EPI sequence (TR=2.2 s, TE=27 ms, flip angle=90°, voxel size=4 mm x 4 mm x 4 mm, 36 slices). In each session, one gradient echo field map sequence was acquired with the same prescription as the functional images. An EyeLink 1000 eye-tracking system (http://www.sr-research.com) allowed continuous monitoring of participants’ eyes in order to check for periods of prolonged eye closure, potentially indicating sleep. Only one participant (P13) demonstrated prolonged eye closures.

#### Task design

##### Motor task design

The motor task was adapted from that used in the Human Connectome Project ^53^. Participants were presented with visual cues that directed them to close and relax their hands, flex and relax their toes, or wiggle their tongue. Each block started with a 2.2 s cue indicating which movement was to be made. After this cue, a centrally-presented caret replaced the instruction and flickered once every 1.1 s (without temporal jittering). Each time the caret flickered, participants executed the proper movement. 12 movements were made per block. Each task run consisted of 2 blocks of each type of movement as well as 3 blocks of resting fixation, which lasted 15.4 s.

##### Mixed block/event-related design task

This task was adapted from experimental conditions reported by ^54^. One task was a spatial coherence discrimination task, which used concentric dot patterns ^74^ that were either 0% or 50% coherent. During this task, participants had to identify each pattern as concentric or random. The other task was a verbal discrimination task. Participants were presented with nouns and verbs and had to identify which type of word was being presented on the screen. Task blocks began with a 2.2 s cue screen indicating which task was to be conducted in the following block. Blocks consisted of 30 trials (half concentric/half nonconcentric for coherence, half noun/half verb for verbal). Stimuli were presented for 0.5 s with a variable 1.7-8.3 s ISI. A stop cue displayed for 2.2 s signaled the end of each task block. Each scan run consisted of two blocks of each task. Task blocks were separated by 44 s periods of rest. For each task, the finger used for each response was counterbalanced within participants across sessions.

### MRI Processing: Dataset 1 & 2

#### Structural Processing

Structural images (T1- and T2-weighted) were corrected for gain field inhomogeneity using FSL Fast ^75^ and aligned to the 711-2B implementation of Talairach atlas space using the 4dfp MRI processing software package (https://readthedocs.org/projects/4dfp/). The 711-2B template conforms to the 1988 Talairach atlas ^76^ according to the method of ^77^. Relative to MNI152, 711-2B space is about 5% smaller and 2° anteriorly rotated about the ear-to-ear axis. Mean T1- and T2-weighted images (T1w and T2w) were computed by coregistration and averaging multiple acquisitions.

Generation of cortical surfaces from the MRI data followed a procedure similar to that previously described in ^45^. First, anatomical surfaces were generated from the participant’s average T1-weighted image in native volumetric space using FreeSurfer’s default recon-all processing pipeline (version 5.3). This pipeline first conducted brain extraction and segmentation. After this step, segmentations were hand-edited to maximize accuracy. Subsequently, the remainder of the recon-all pipeline was conducted on the hand-edited segmentations, including generation of white matter and pial surfaces, inflation of the surfaces to a sphere, and surface shape-based spherical registration of the participant’s original surface to the fsaverage surface ^78,79^. The fsaverage-registered left and right hemisphere surfaces were brought into register with each other using deformation maps from a landmark-based registration of left and right fsaverage surfaces to a hybrid left-right fsaverage surface (‘fs_LR’; ^80^). These fs_LR spherical template meshes were input to a flexible Multi-modal Surface Matching (MSM) algorithm using sulcal features to register templates to the atlas mesh ^81^. These newly registered surfaces were then down-sampled to a 32,492 vertex surface (fs_LR 32k) for each hemisphere. These various surfaces in native stereotaxic space were then transformed into atlas space (711-2B) by applying the previously calculated T1-to-atlas transformation.

#### fMRI Preprocessing

Functional data were preprocessed to reduce artifacts and to maximize cross-session registration. All sessions underwent correction of odd vs. even slice intensity differences attributable to interleaved acquisition, intensity normalization to a whole brain mode value of 1000, and within run correction for head movement. Atlas transformation was computed by registering the mean intensity image from a single BOLD session to atlas space via the average high-resolution T2-weighted image and average high-resolution T1-weighted image. All subsequent BOLD sessions were linearly registered to this first session. This atlas transformation, mean field distortion correction (see below), and resampling to 3-mm isotropic atlas space were combined into a single interpolation using FSL’s applywarp tool. All subsequent operations were performed on the atlas-transformed volumetric time series.

#### Distortion Correction

A mean field map was generated based on the field maps collected in each participant ^43^. This mean field map was then linearly registered to each session and applied to that session for distortion correction. To generate the mean field map the following procedure was used: (1) 4 Field map magnitude images were mutually co-registered. (2) Transforms between all sessions were resolved. Transform resolution reconstructs the n-1 transforms between all images using the n(n-1)/2 computed transform pairs. (3) The resolved transforms were applied to generate a mean magnitude image. (4) The mean magnitude image was registered to an atlas representative template. (5) Individual session magnitude image to atlas space transforms were computed by composing the session-to-mean and mean-to-atlas transforms. (6) Phase images were then transformed to atlas space using the composed transforms, and a mean phase image in atlas space was computed. Application of mean field map to individual fMRI sessions: (1) For each session, field map uncorrected data were registered to atlas space, as above. (2) The generated transformation matrix was then inverted and applied to the mean field map to bring the mean field map into the session space. (3) The mean field map was used to correct distortion in each native-space run of resting state and task data in the session. (4) The undistorted data were then re-registered to atlas space. (5) This new transformation matrix and the mean field map then were applied together to resample each run of resting state and task data in the session to undistorted atlas space in a single step.

#### RSFC Preprocessing

Additional preprocessing steps to reduce spurious variance unlikely to reflect neuronal activity were executed as recommended in ^82,83^. First, temporal masks were created to flag motion-contaminated frames. Motion estimate time courses were filtered retain effects occurring below 0.1 Hz in order to eliminate “pseudomotion” induced by breathing-related motion of the chest altering the B0 field ^84^. Motion contaminated volumes were then identified by frame-by-frame displacement (FD). Frames with FD > 0.2mm were flagged as motion-contaminated.

After computing the temporal masks for high motion frame censoring, the data were processed with the following steps: (i) demeaning and detrending, (ii) linear interpolation across censored frames using so that continuous data can be passed through (iii) a band-pass filter (0.005 Hz < f < 0.01 Hz) without re-introducing nuisance signals ^85^ or contaminating frames near high motion frames.

Next, the filtered BOLD time series underwent a component-based nuisance regression approach ^45^. Nuisance regression using time series extracted from white matter and cerebrospinal fluid (CSF) assumes that variance in such regions is unlikely to reflect neural activity. Variance in these regions is known to correspond largely to physiological noise (e.g., CSF pulsations), arterial pCO2-dependent changes in T2*-weighted intensity and motion artifact; this spurious variance is widely shared with regions of interest in gray matter. We also included the mean signal averaged over the whole brain as a nuisance regressor. Global signal regression (GSR) has been controversial. However, the available evidence indicates that GSR is a highly effective de-noising strategy ^82,86^.

Nuisance regressors were extracted from white matter and ventricle masks, first segmented by FreeSurfer ^87^, then spatially resampled in register with the fMRI data. Voxels surrounding the edge of the brain are particularly susceptible to motion artifacts and CSF pulsations ^88,89^; hence, a third nuisance mask was created for the extra-axial compartment by thresholding the temporal standard deviation image (SDt > 2.5%), excluding a dilated whole brain mask. Voxelwise nuisance time series were dimensionality reduced as in CompCor ^90^, except that the number of retained regressors, rather than being a fixed quantity, was determined, for each noise compartment, by orthogonalization of the covariance matrix and retaining components 5 ordered by decreasing eigenvalue up to a condition number of 30 (max eigenvalue / min eigenvalue > 30). The retained components across all compartments formed the columns of a design matrix, X, along with the global signal, its first derivative, and the six time series derived by retrospective motion correction. The columns of X are likely to exhibit substantial co-linearity. Therefore, to prevent numerical instability owing to rank-deficiency during nuisance regression, a second-level SVD was applied to XXT to impose an upper limit of 250 on the condition number. This final set of regressors was applied in a single step to the filtered, interpolated BOLD time series, with censored data ignored during beta estimation. Censored frames were then excised from the data for all subsequent analyses.

#### Surface processing and CIFTI generation of BOLD data

Surface processing of BOLD data proceeded through the following steps. First, the BOLD fMRI volumetric timeseries (both resting-state and task) were sampled to each participant’s original mid-thickness left and right-hemisphere surfaces (generated as the average of the white and pial surfaces) using the ribbon-constrained sampling procedure available in Connectome Workbench 1.0 ^91^. This procedure samples data from voxels within the gray matter ribbon (i.e., between the white and pial surfaces) that lie in a cylinder orthogonal to the local mid-thickness surface weighted by the extent to which the voxel falls within the ribbon. voxels with a timeseries coefficient of variation 0.5 standard deviations higher than the mean coefficient of variation of nearby voxels (within a 5 mm sigma Gaussian neighborhood) were excluded from the volume to surface sampling, as described in ^92^. Once sampled to the surface, timecourses were deformed and resampled from the individual’s original surface to the 32k fs_LR surface in a single step using the deformation map generated above (in “Cortical surface generation”). This resampling allows point-to-point comparison between each individual registered to this surface space.

These surfaces were then combined with volumetric subcortical and cerebellar data into the CIFTI format using Connectome Workbench, creating full brain timecourses excluding nongray matter tissue. Subcortical (including accumbens, amygdala, caudate, hippocampus, pallidum, putamen, and thalamus) and cerebellar voxels were selected based on the FreeSurfer segmentation of the individual participant’s native-space average T1, transformed into atlas space, and manually inspected. Finally, the BOLD timecourses were smoothed with a geodesic 2D (for surface data) or Euclidean 3D (for volumetric data) Gaussian kernel of σ = 2.55 mm.

#### Regression of adjacent cortical tissue from RSFC BOLD

Many subcortical areas, such as dorsal cerebellum and lateral putamen, are in close anatomical proximity to cortex, resulting in spurious functional coupling between the cortical vertices and adjacent subcortical voxels. To reduce this artifact, RSFC BOLD time series from all vertices falling within 20mm Euclidean distance of a source voxel were averaged and then regressed from the voxel time series ^45,73,93,94^. The resulting residual timeseries were used for all subsequent analyses.

### Dataset 3: Multi-echo dataset

#### Participants

Data were collected from 4 healthy, right-handed adults (0 female; ages 29, 39, 24, and 31; identified here as P04-P07). Other findings using these participants have been previously reported in ^21,95^. Informed consent was obtained from all participants. The study was approved by the Weill Cornell School of Medicine Institutional Review Board.

#### MRI Acquisition

Data were acquired on a Siemens Magnetom Prisma 3T scanner at the Citigroup Biomedical Imaging Center of Weill Cornell’s medical campus using a Siemens 32-channel head coil. Multi-echo, multi-band resting-state fMRI scans were collected using a T2*-weighted echo-planar sequence covering the full brain (TR: 1355 ms; TE1: 13.40 ms, TE2: 31.11 ms, TE3: 48.82 ms, TE4: 66.53 ms, and TE5: 84.24 ms; FOV: 216 mm; flip angle: 68; 2.4mm isotropic; 72 slices; AP phase encoding direction; in-plane acceleration factor: 2; and multi-band acceleration factor: 6) with 640 volumes acquired per scan for a total acquisition time of 14 min and 27 s. This sequence was generously provided by the Center for Magnetic Resonance Research (CMRR) at the University of Minnesota. A pair of spin echo EPI images with opposite phase encoding directions (AP and PA) but identical geometrical parameters and echo spacing were acquired to correct spatial distortions. High-resolution (MPRAGE) T1-weighted image (TR: 2400 ms; TE: 2.28 ms; FOV: 256; flip angle: 90, and 208 sagittal slices with a 0.8 mm thickness) and T2-weighted anatomical images (TR: 3200 ms; TE: 563 ms; FOV: 256; flip angle: 8, and 208 sagittal slices with a 0.8 mm thickness) were acquired. Custom headcases were obtained from Caseforge (https://ipira.berkeley.edu/caseforge-inc) for each participant to improve comfort and minimize head motion during scanning ^96^.

### MRI Processing: Dataset 3

#### Cortical Surface Generation

The average T1- and T2-weighted images were cropped to a smaller field of view (170mm in z plane), co-registered using FSL’s epi_reg tool (via a boundary-based cost function with 6 DOF), and corrected for intensity inhomogeneities ^97^. The T1- and T2-weighted images were co-registered to an MNI atlas (hereafter referred to as “ACPC” alignment) using a rigid 6 DOF FLIRT transformation. Cortical surfaces were generated using Freesurfer’s “recon-all.v6.hires” pipeline. Pial surface placement was refined using the co-registered T2-weighted image by specifying the “-T2pial” option. Midthickness surfaces were obtained by averaging the pial and white surfaces. Fsaverage-registered left and right hemisphere surfaces (pial, white, and midthickness) were brought into register with each other in fs_LR space ^80^ and resampled to the computationally tractable resolution of 32k vertices using Connectome Workbench command line utilities.

#### fMRI Preprocessing

Preprocessing of multi-echo data minimized spatial interpolation and volumetric smoothing while preserving the alignment of echoes. The single-band reference (SBR) images (five total; one per echo) for each scan were averaged. The resultant average SBR images were aligned, averaged, co-registered to the ACPC aligned T1-weighted anatomical image, and simultaneously corrected for spatial distortions using FSL’s topup and epi_reg programs. Freesurfer’s bbregister algorithm ^98^ was used to refine this co-registration. For each scan, echoes were combined at each time point and a unique 6 DOF registration (one per volume) to the average SBR image was estimated using FSL’s MCFLIRT tool ^99^ using a 4-stage (sinc) optimization. All of these steps (co-registration to the average SBR image, ACPC alignment, and correcting for spatial distortions) were concatenated using FSL’s convertwarp tool and applied as a single spline warp to individual volumes of each echo after correcting for slice time differences using FSL’s slicetimer program. All denoising was performed on these preprocessed, ACPC-aligned images.

#### Multi-echo denoising

Multi-echo ICA (ME-ICA; ^100,101^ denoising designed to isolate spatially structured T2∗-(neurobiological; “BOLD-like”) and S0-dependent (non-neurobiological; “not BOLD-like”) signals was performed using a modified version of the “tedana.py” workflow (https://tedana.readthedocs.io/en/latest/). In short, the preprocessed, ACPC-aligned echoes were first combined according to the average rate of T2∗ decay at each voxel across all time points by fitting the monoexponential decay, S(t) = S0e -t / T2∗, using the “nlinfit.m” function in MATLAB with least-squares optimization and the initial coefficient values obtained from a linear model fit to the log of the data. From these T2∗ values, an optimally combined multi-echo (OC-ME) time-series was obtained by combining echoes using a weighted average (WTE = TE ∗ e -TE/ T2∗), as in ^102^. The covariance structure of all voxel time-courses was used to identify major signals in the resultant OC-ME time-series using principal component and independent component analysis. Components were classified as either T2∗-dependent (and retained) or S0-dependent (and discarded), primarily according to their decay properties across echoes following the decision tree described in ^100^. Mean gray matter time-series regression was subsequently performed to remove spatially diffuse noise. Temporal masks were generated for censoring high motion time-points using a frame-wise displacement (FD; ^103^ threshold of 0.3 mm and a backward difference of two TRs (2 ∗ 1.355 = 2.77 s), for an effective sampling rate comparable to historical FD measurements (approximately 2 to 4 s; ^96^. Prior to the FD calculation, head realignment parameters were filtered using a stopband Butterworth filter (0.2 - 0.35 Hz) to attenuate the influence of respiration ^96^.

#### Surface processing and CIFTI generation of BOLD Data

The denoised fMRI time-series was mapped to the midthickness surfaces (using the “-ribbon-constrained” method), combined into the Connectivity Informatics Technology Initiative (CIFTI) format, and spatially smoothed with geodesic (for surface data) and Euclidean (for volumetric data) Gaussian kernels (σ = 2.55 mm) using Connectome Workbench command line utilities (Glasser et al., 2013). Signals were normalized (z-scored). This yielded time courses representative of the entire cortical surface, subcortex (accumbens, amygdala, caudate, hippocampus, pallidum, putamen, and thalamus), and cerebellum, but excluding non-gray matter tissue. Signals from adjacent cortex were regressed from subcortical voxels, as in Datasets 1 and 3.

### Analysis

#### Mapping Subnetwork Structure

The network organization of each participant’s brain was delineated following ^35^ using the graph-theory based Infomap algorithm for community detection ^104^. In this approach, we calculated the cross-correlation matrix of the time courses from all brain vertices (on the cortical surfaces) and voxels (in subcortical structures), concatenated across sessions. Correlations between vertices/voxels within 30 mm of each other were set to zero in this matrix to avoid basing network membership on correlations attributable to spatial smoothing. Geodesic distance was used for within-hemisphere surface connections and Euclidean distance for subcortical-to-cortical connections. Connections between subcortical structures were disallowed, as we observed extremely high correlation values within nearly the entire basal ganglia that would prevent network structures from emerging. Interhemispheric connections between the cortical surfaces were retained, as smoothing was not performed across the midsagittal plane.

We observed that connectivity patterns within regions known to have low BOLD signal due to susceptibility artifact dropout (e.g., ventral anterior temporal lobe and portions of orbitofrontal cortex) were unstructured and inconsistent across individuals. To avoid having the delineated network structures distorted by regions with poor signal, connections to regions with average mode-1000 normalized BOLD signal <750 were set to zero (as in ^105,106^).

The cross-correlation matrix was then thresholded to retain at least the strongest 0.1% of connections to each vertex and voxel ^21,35,49^. Note that this thresholding approach differs from previous approaches for forming brain graphs from functional connectivity data (e.g., ^2,5^). The typical procedure applies a uniform edge density threshold to all functional connectivity values in the brain. The weakness of the uniform threshold approach is that subcortical structures generally have decreased BOLD signal-to-noise relative to cortex due to their greater distance from the MR head coil. The result is that functional connectivity patterns seeded from striatal voxels have weaker peak connectivity strengths, even though they may appear well organized and coherent with known cortical networks. Thus, with a uniform threshold, these regions are frequently not identified as being networked with cortical regions.

By contrast, the current approach thresholds the connectivity maps seeded from each cortical or subcortical point in the brain separately, always retaining at least the 0.1% strongest connections. Previous validation of this procedure ^35^ has shown that the 0.1% density threshold identifies subnetwork divisions in resting data that best explain task activations in these participants.

The thresholded matrices were used as inputs for the Infomap algorithm, which calculated community assignments separately for each threshold. The resulting communities represent subnetworks in the brain. Small networks with 10 or fewer vertices/voxels were considered unassigned and removed from further consideration. The above analysis was conducted in each individual participant.

#### Identifying Matched CON Subnetworks in Individuals

For each participant, we considered only subnetworks that had at least some representation within the traditional distribution of the cingulo-opercular network including the insula, anterior cingulate cortex extending dorsally into dorsomedial prefrontal cortex, and supermarginal gyrus ^2,5,6,8–10^. Following ^35^, we visually examined the cortical and subcortical topographies of each subnetwork with CON representation, as well as their topological arrangement relative to each other. After careful consideration of the subnetworks observable in this population, we initially identified four potential discrete CON subnetworks. However, we observed that 1) one of the subnetworks (in middle insula and dorsal postcentral gyrus) could not be separately delineated in all participants; and 2) that same subnetwork was not dissociable from the Central subnetwork in its connectivity, lag ordering, or task responses. See Supplementary Figure S3 for details. Thus, we concluded that the CON was best described by three discrete subnetworks. These matched subnetworks were identified based on the following heuristic rules:

The Anterior (Decision) subnetwork was present bilaterally with representations in the dorsal anterior cingulate, anterior insula, and anterior prefrontal cortex.

The Central (Action) subnetwork was present posterior to the Anterior subnetwork in the more posterior aspect of the dorsal anterior cingulate and extended into posterior dorsal medial prefrontal cortex, as well as in middle insula, and anterior superior parietal cortex.

The Lateral (Feedback) subnetwork was present in the supermarginal gyrus, posterior anterior insula, anterior and posterior inferior frontal gyrus, and anterior prefrontal cortex.

After identifying subnetworks on the cortex, we examined subcortical areas (basal ganglia, thalamus, and cerebellum) for representation of our selected subnetworks in these regions.

#### Visualizing Subnetwork Overlap across Participants

For each matched subnetwork, the number of individuals with each subnetwork present was calculated for each cortical vertex. For subcortical voxels, some participants (Plasticity and MSC) were in Talaraich space. To represent overlap, we first transformed each of these participants into MNI space. Overlap was then calculated across MNI-space participant subnetworks as the number of individuals with the subnetwork present in each MNI-space voxel. This procedure produced maps of the density of each subnetwork across participants.

#### Meta-Analytic Network Annotation analysis

We investigated functions associated with each subnetwork by leveraging text-mining and meta-analyses techniques in the brain mapping Neurosynth database (https://github.com/neurosynth/neurosynth-data). Neurosynth compiles neuroimaging studies and generates probabilistic mappings based on article term frequencies and term-to-activation correlations ^50^. Neurosynth functionality natively includes mapping single [X, Y, Z] coordinates to meta-analytic descriptor terms common in the neuroimaging literature. This is accomplished by automatically text-mining both activity peaks and descriptor terms from a huge corpus of neuroimaging papers. Then, papers can be identified in which a reported activity peak falls within a short distance of the target coordinate. At the time the Neurosynth database was downloaded, it contained over 500,000 activation peaks from over 14,000 fMRI papers. Each paper is labeled with at least one term automatically mined from its text, and each term has a weighting for each paper reflecting the prevalence of that term in the text.

Here, we expanded this functionality by mapping not just a single coordinate, but instead simultaneously mapping the multiple regions within each subnetwork. We first identified the congruent clusters of each subnetwork present in the group. This was accomplished by thresholding the subnetwork density maps (representing cross-participant overlap, described above) to retain all cortical vertices in which >10% of participants had the subnetwork. For each subnetwork, we then identified all studies within the Neurosynth database with a collection of activation peaks that “matched” the subnetwork. Specifically, the study had to report an activation peak <2mm from at least 30% of subnetwork regions. Thus, a given study must have elicited activity near a substantial number of the subnetwork’s regions in order to match that subnetwork. Varying these parameters did not substantially alter findings reported here.

All terms associated with matching studies that were related to mental or task-related function (e.g., “effort”, “oddball”, “language”, “delay”, “covert”), were retained, while all terms related to aspects of the participant population (e.g. “male”), brain location (e.g. “occipital”, “network”), or anything unrelated (e.g. “voxel”, “peripheral”, “extra”) were excluded. This restricted the database to 742 terms.

Terms in the Neurosynth database have “weights” for each study ranging from 0 to 1, indicating the prevalence of that term within the text-mined paper. For each term, we tested whether that term’s weights differed significantly across different subnetworks. Specifically, the weights found for that term in each study matched to each subnetwork were all entered into a one-way ANOVA testing for effects of the identity of the matching subnetwork. Significance in this test indicates significant differences in the weight of that term across subnetworks. If the ANOVA was significant, that term was associated with the subnetwork exhibiting the largest average weight. Significance was tested both at p<.05, FDR-corrected for the number of terms tested, as well as (for exploratory purposes) at p<.05 uncorrected.

#### Mapping Individual-Specific Large-Scale Networks

We identified the set of canonical large-scale networks using the individual-specific network matching approach described in ^2^. Briefly, the Infomap algorithm was applied to each participant’s correlation matrix thresholded at a range of edge densities spanning from 0.01% to 5%. At each threshold, the algorithm returned community identities for each vertex and voxel. Communities were labeled by matching them at each threshold to a set of independent group average networks described in ^2^. The matching approach proceeded as follows: 1) At each density threshold, all identified communities were compared with the independent group networks using the Jaccard Index of spatial overlap. 2) The community with the best match (highest overlap) to one of the independent networks was assigned that network identity, and then not considered for further comparison with other independent networks within that threshold. Matches lower than Jaccard = 0.1 were not considered (to avoid matching based on only a few vertices). Matches were first made with the large, well-known networks (in order: default, lateral visual, motor hand, motor mouth, frontoparietal, and dorsal attention, language), and then to the smaller networks (salience, parietal memory, contextual association, medial visual, motor foot, and somato-cognitive action). In each individual and in the average, a “consensus” network assignment was derived by collapsing assignments across thresholds, giving each node the assignment it had at the sparsest possible threshold at which it was successfully assigned to one of the known group networks.

#### Calculating Connectivity to Other Networks / Subnetwork-Network Relationships

For each CON subnetwork matched across participants, we calculated the average fMRI time course across subnetwork voxels/vertices. We then calculated the average time course across voxels/vertices of the large-scale Default Mode, Visual, Fronto-Parietal, Dorsal Attention, Language, Salience, Somatomotor Hand, Somatomotor Face, Somatomotor Foot, Auditory, Parietal Memory, Context, and Somato-Cognitive Action networks. Voxels/vertices that overlapped with that subnetwork were excluded from these averages. We then calculated the functional connectivity between each CON subnetwork and each of the 12 large-scale networks as the Fisher-transformed correlation of the two-time courses.

For each large-scale network, we compared the connectivity strengths of the subnetworks against each other using paired *t* tests (subnetwork vs. subnetwork) and applying FDR correction for multiple comparisons across all network / pairwise subnetwork tests.

We calculated pairwise functional connectivity strengths between the CON subnetworks themselves to better understand the integration of these subnetworks within the larger-scale CON. We directly compared connectivity between different pairs of CON subnetworks using paired t-tests.

Visualization of subnetwork-network relationships in individual participants was conducted using spring-embedded plots ^5^, as implemented in Gephi (https://gephi.org/). In each participant, nodes were defined as contiguous cortical subnetwork or network clusters larger than 20 mm^2^ from each matched CON subnetworks, as well as from within the other large-scale networks shown to be strongly connected to CON subnetworks. Pairwise connectivity between nodes was calculated as the Z-transformed correlation of their mean time courses. For visualization purposes, graphs were constructed by thresholding node-to-node connectivity matrices at 15% density. These graphs were then imported into Gephi. For three participants (P05, P07, and P13), whole networks (always either Salience or Dorsal Attention) were disconnected from the rest of the graph at this density threshold. Since the goal was to visualize connectivity with these networks, in these cases we systematically increased the density threshold in 5% increments until all networks were connected to the graph. For these three participants, the graph became connected and was visualized at 45%, 30%, and 20% densities, respectively.

#### Calculating Subnetwork Time Delays

We computed Time Delay (TD) estimates using an adaptation of a previously published method ^107^. Briefly, the method of ^107^ computes a lagged cross-covariance function (CCF) between timecourses for each session. We observed that these cross-covariance functions were more stable and consistent across participants when they were restricted to timepoints with large signal fluctuations in the regions of interest. This observations follows recent work suggesting that large-amplitude fluctuations allow more precise estimation of individual-specific functional connectivity patterns^108–110^. Thus, we restricted timepoints of interest to those in which at least 0.1% of the vertices within the CON subnetworks exhibited a large signal fluctuation (> 5 standard deviations above the vertex mean), as well as six seconds before and after that large fluctuation. For each of these time segments, we computed the lagged CCF between each pair of cortical vertex time courses. To account for censored frames, we computed CCFs over blocks of contiguous frames and averaged these CCFs, weighted by block duration, to obtain a single segment CCF. We excluded TDs greater than 4 s as, in our experience, these tend to reflect sampling error or artifact. Thus, CCFs were computed over three TR shifts in the positive and negative directions, making the minimum block duration [3 (TR shifts) + 1 (zero-lag)] x TR = 8.8 s (Dataset 1 & 2); 5.42 s (Dataset 3). Lags were then more precisely determined by estimated the cross-covariance extremum of the session CCF using three-point parabolic interpolation. The resulting lags were assembled into an antisymmetric matrix capturing all pairwise TDs (TD matrix) for each time segment, which was averaged across time segments to yield participant-level TD vertexwise matrices.

Each participant’s TD matrix was averaged across rows to summarize the average time shift from each vertex to all other vertices. These average time shifts were then averaged across vertices for each subnetwork and network of interest. A one-way ANOVA tested whether there were any differences in row-average TD across the subnetworks, as well as the SCAN and the Somatomotor Hand network. Post-hoc t-tests determined the relative temporal ordering of networks.

#### Task Analysis

Task evoked activations were modeled individually for each vertex and voxel with a general linear model (GLM) ^111^, using in-house image analysis software written in IDL (Research Systems, Inc.). First level GLM analyses were conducted separately for each session in a given participant, and second level within-participant analyses were run on the session-wise beta values of a single participant. Planned second-level contrasts were evaluated as paired voxel/vertex-wise t-tests comparing these beta values, and the resulting t-values in each voxel/vertex were then Z-transformed for further analysis.

We included tasks that had two different types of designs (motor = block design, spatial/verbal discrimination = mixed block/event-related design). In the block design motor task, a block regressor was convolved with a canonical hemodynamic response to model the five experimental conditions: tongue, left hand, right hand, left foot, right foot.

The spatial and verbal discrimination tasks were jointly modeled in a mixed block-event related design. Events were modeled with an FIR model (as above, with 8 timepoints for each event); separate event regressors were included for the start and end cues in each task, and for the different trial types (noun, verb, 50% coherence, 0% coherence). The block (sustained activity) was modeled with a square block regressor, with separate regressors for sustained activity in the semantic and coherence task. Given the low number of error trials, errors were not modeled in any task. In addition to these terms, constant and linear effects were modeled for each run.

The task contrasts of interest included: 1) Tongue > baseline, 2) Left Hand > baseline, 3) Right Hand > baseline, 4) Left Leg > baseline, 5) Right Leg > baseline, 6) Spatial Discrimination Task > baseline, 7) Verbal Discrimination Task > baseline. We calculated the average activation (z-scores) for each subnetwork in the vertices involved in each of the tasks.

