## Supplemental Material for "Substructure of the brain’s Cingulo-Opercular network"

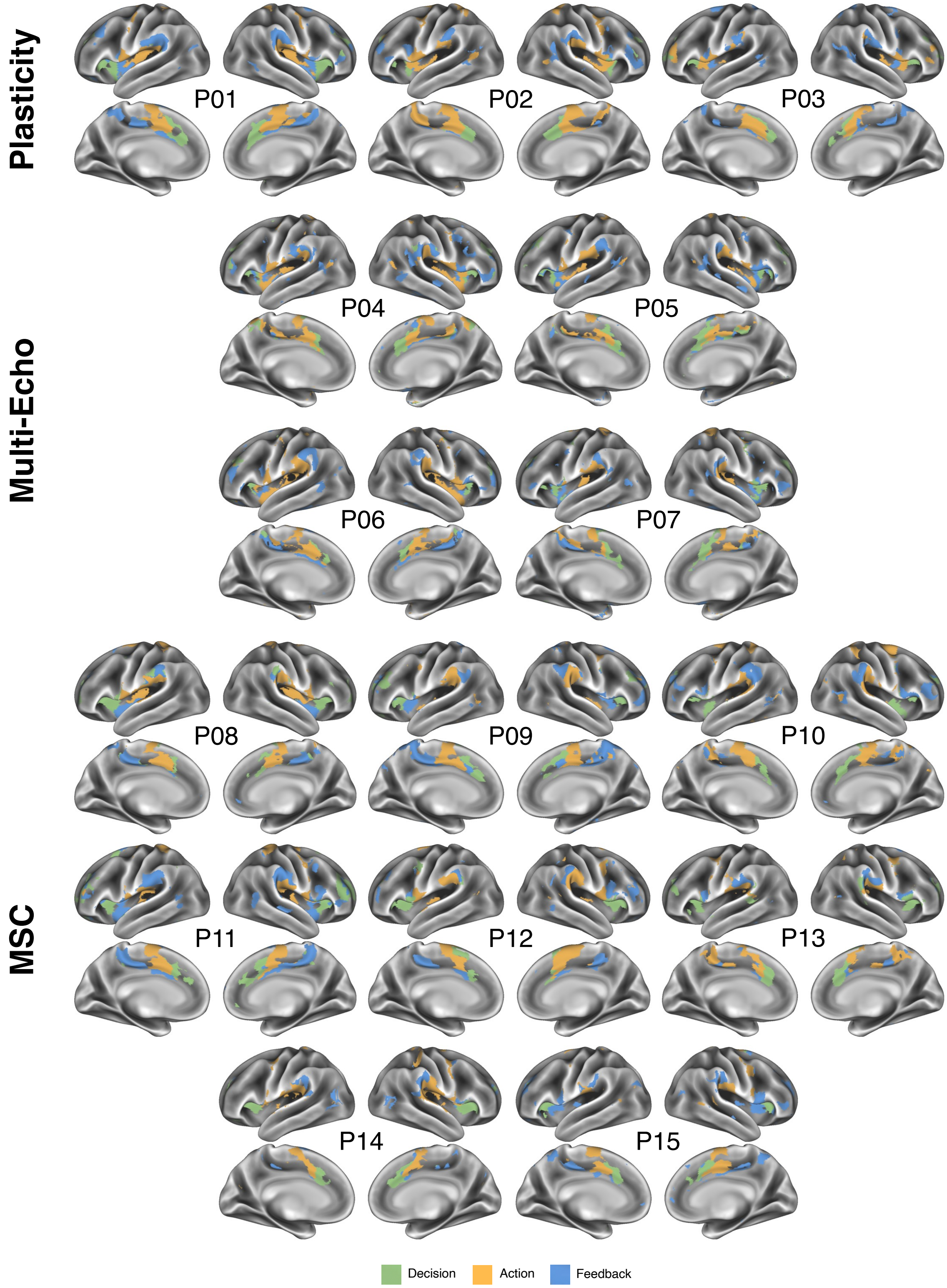


**Figure S1: Individual CON Subnetwork Maps.** For each participant, the map illustrates the topography of the three identified CON subnetworks.

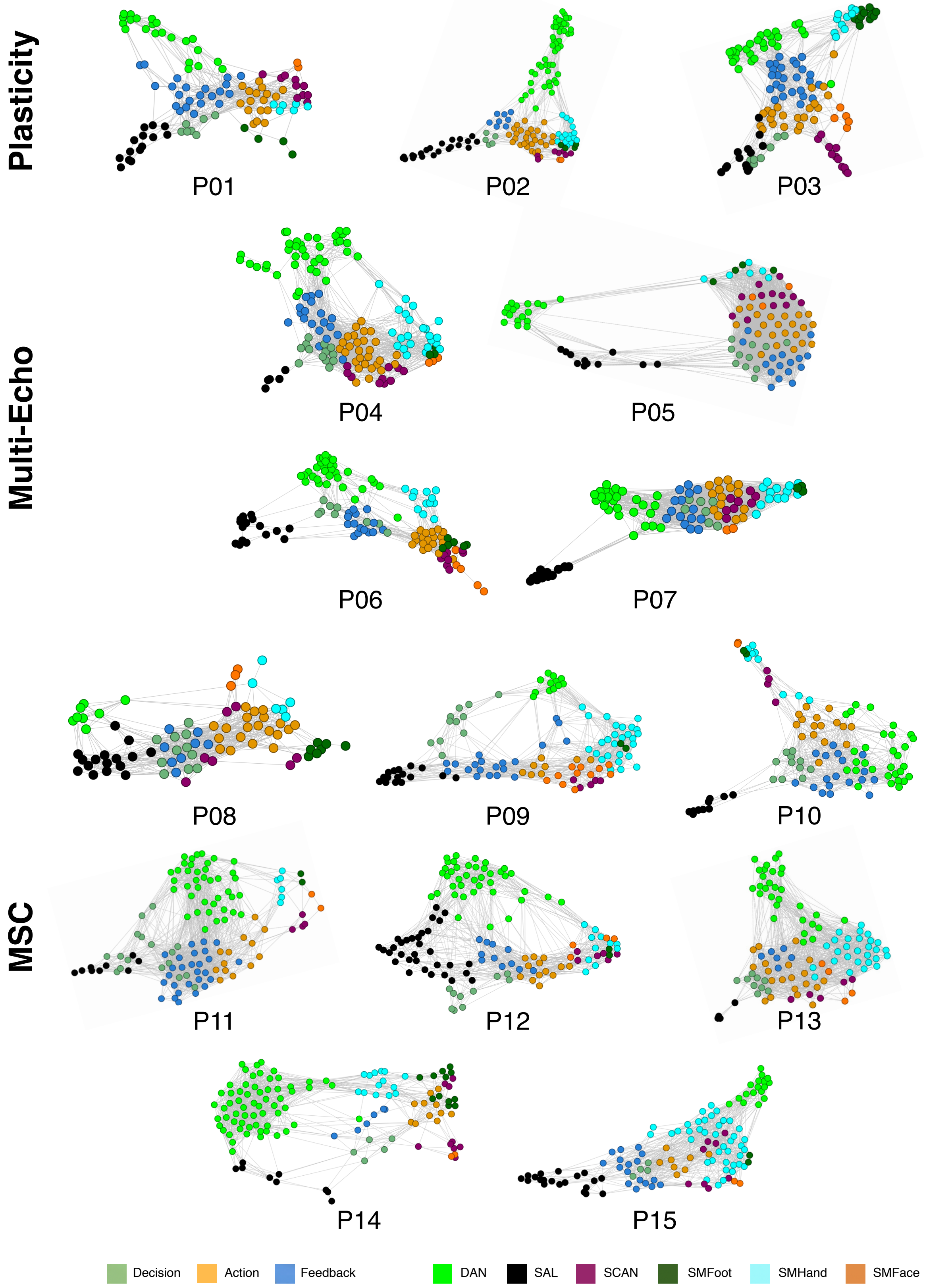


**Figure S2: Individual CON subnetwork spring embedding plots.** For each participant, the plot illustrates connections between CON Subnetworks and the Salience, DAN, SCAN, and somatomotor foot, hand, and face networks.

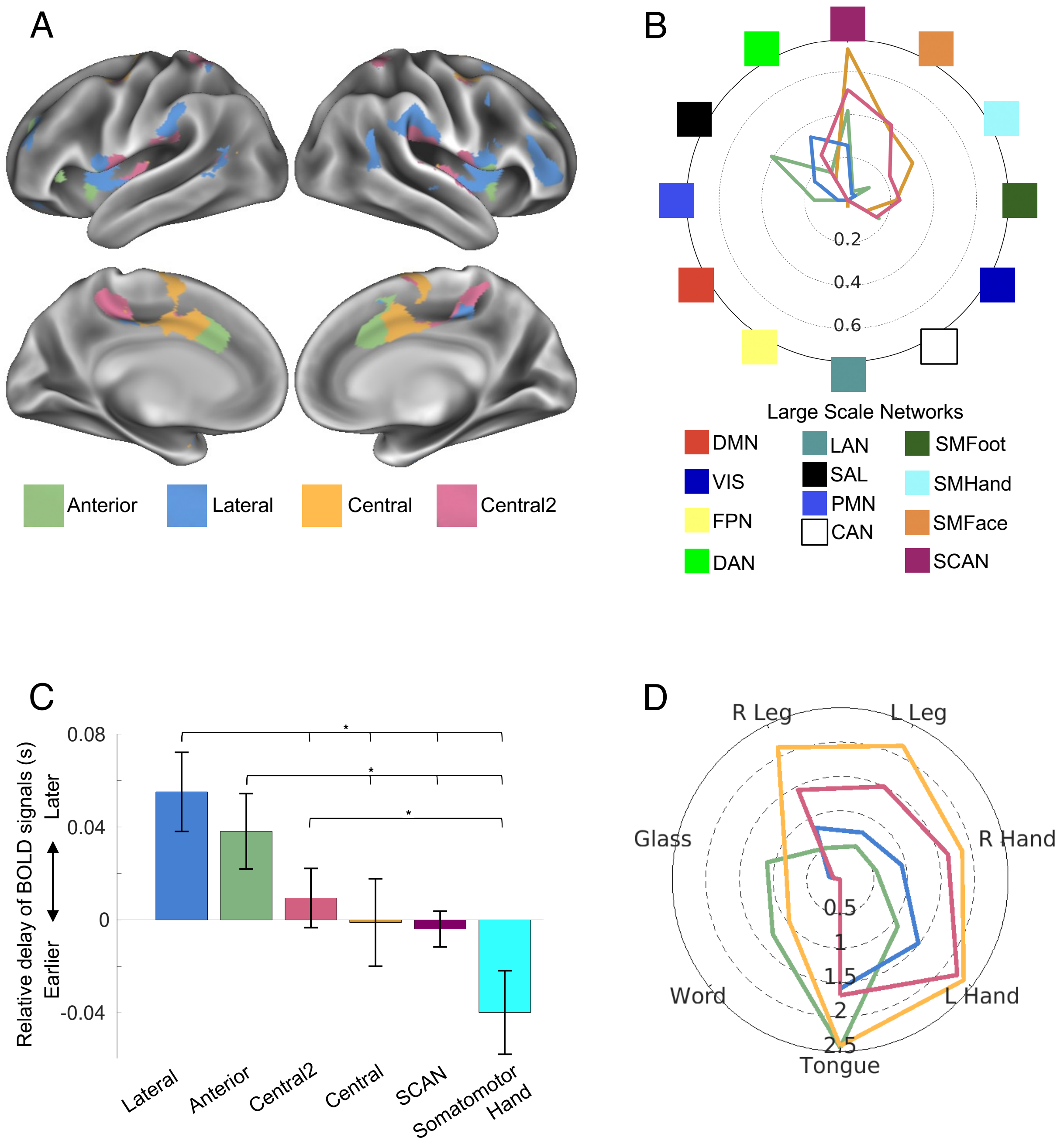


**Figure S3: Analyses conducted with four CON subnetworks.** A) Four CON subnetworks in an example subject. The Central subnetwork could be divided into two discrete subnetworks in some but not all subjects. B) Functional connectivity of four CON subnetworks to large-scale networks. The radial axis indicates the strength of functional connectivity Z(r) between each CON subnetwork and each large-scale network. Negative connectivity values are not represented. Colors of large-scale networks are shown at the bottom. The two Central subnetworks exhibited very similar connectivity. C) Temporal ordering of signals within four CON subnetworks, and in SCAN and Somatomotor Hand networks. Standard error across subjects is indicated by error bars. A one-way ANOVA indicated a significant main effect of subnetwork/network identity ($p = 0.0005$). * indicates $p < 0.05$ for post-hoc paired t-tests. The two Central subnetworks exhibited very similar ordering of signals. D) Task activation of four CON subnetworks in motor, spatial discrimination, and verbal discrimination tasks. The radial axis indicates the z-score compared to a baseline condition averaged across all subjects and across all vertices in each subnetwork. The two Central subnetworks exhibited very similar activation profiles.

**Supplementary Table 1: CON Subnetwork vs Large Scale Network Connectivity.** Paired t-tests compared functional connectivity to a large-scale network between two different CON subnetworks. Asterisks indicate significant differences (FDR corrected to alpha < 0.05) between pairs of subnetworks.

| | Subnetwork comparison | | |
|---|---|---|---|
| Connectivity to: | Decision vs Action | Decision vs Feedback | Action vs Feedback |
| Default Mode | t(14) = -3.2; p = 0.006* | t(14) = -1.2; p = 0.26 | t(14) = 3.1; p = 0.008* |
| SCAN | t(14) = -5.9; p < 0.001* | t(14) = 3.3; p = 0.006* | t(14) = 8.4; p < 0.001* |
| Visual | t(14) = 0.02; p = 0.98 | t(14) = 2.1; p = 0.06 | t(14) = 2.1; p = 0.05 |
| Fronto-Parietal | t(14) = 4.4; p < 0.001* | t(14) = 1.0; p = 0.31 | t(14) = -4.2; p < 0.001* |
| Dorsal Attention | t(14) = -0.4; p = 0.72 | t(14) = -3.6; p = 0.003* | t(14) = -3.2; p = 0.006* |
| Language | t(14) = 1.5; p = 0.15 | t(14) = 3.2; p = 0.007* | t(14) = 2.3; p = 0.04 |
| Salience | t(14) = 5.6; p < 0.001* | t(14) = 2.9; p = 0.01* | t(14) = -3.6; p = 0.003* |
| Somatomotor Hand | t(14) = -9.3; p < 0.001* | t(14) = -0.5; p = 0.65 | t(14) = 6.2; p < 0.001* |
| Somatomotor Face | t(14) = -5.8; p < 0.001* | t(14) = 1.2; p = 0.24 | t(14) = 6.9; p < 0.001* |
| Somatomotor Foot | t(14) = -6.4; p < 0.001* | t(14) = -0.2; p = 0.85 | t(14) = 5.5; p = 0.001* |
| Parietal Memory | t(14) = 7.0; p < 0.001* | t(14) = 1.8; p = 0.09 | t(14) = -2.2; p = 0.04 |
| Context | t(14) = 0.9; p = 0.38 | t(14) = -0.8; p = 0.47 | t(14) = -2.3; p = 0.04 |

**Supplementary Table 2: Temporal ordering of signals.** T-tests describing differences in temporal ordering between pairs of networks and subnetworks.

| | Feedback | Decision | Action | SCAN |
|---|---|---|---|---|
| Decision | t(14) = 1.4; p = 0.17 | | | |
| Action | t(14) = 2.8; p = 0.01 | t(14) = 1.3 p = 0.21 | | |
| SCAN | t(14) = 2.6; p = 0.01 | t(14) = 1.1; p = 0.29 | t(14) = -0.4; p = 0.69 | |
| Somatomotor Hand | t(14) = 3.5; p = 0.001 | t(14) = 2.4; p = 0.02 | t(14) = 1.7; p = 0.09 | t(14) = 2.0; p = 0.05 |

**Supplementary Table 3: MSC Task Connectivity.** Paired t-tests compared activation vs baseline in each task between two different CON subnetworks. Asterisks indicate significant differences in activation between two subnetworks (FDR-corrected to alpha < 0.05) for a specific task condition.

| | Subnetwork comparison | | |
|---|---|---|---|
| Task: | Decision vs Action | Decision vs Feedback | Action vs Feedback |
| Tongue | t(9) = 1.3; p = 0.21 | t(9) = 4.0; p = 0.003* | t(9) = 3.5; p = 0.006* |
| Left Hand | t(9) = -8.1; p < 0.001* | t(9) = -1.4; p = 0.20 | t(9) = 4.7; p = 0.001* |
| Right Hand | t(9) = -5.1; p < 0.001 | t(9) = -0.8; p = 0.43 | t(9) = 3.8; p = 0.004* |
| Left Foot | t(9) = -7.3; p < 0.001* | t(9) = -0.7; p = 0.52 | t(9) = 4.7; p = 0.001* |
| Right Foot | t(9) = -4.9; p < 0.001 | t(9) = -0.7; p = 0.51 | t(9) = 4.4; p = 0.002* |
| Spatial Discrimination | t(9) = 4.4; p = 0.002* | t(9) = 5.7; p = 0.003* | t(9) = 3.1; p = 0.01* |
| Verbal Discrimination | t(9) = 6.4; p = 0.001* | t(9) = 7.3; p < 0.001* | t(9) = 4.2; p = 0.002* |
